# Geographic distribution of the Birmingham Darter *Etheostoma birminghamense*

**DOI:** 10.1101/2025.11.05.686853

**Authors:** Chase D. Brownstein, Konstantinos Andriotis, Christopher B. Haynes, Nathaniel D. Sturm, Bernard R. Kuhajda, Thomas J. Near

## Abstract

Southeastern North America harbors the richest freshwater biodiversity hotspot in the northern hemisphere and is home to numerous species with extremely narrow ranges. Among these are the six species of the *Etheostoma chermocki* species complex, which exclusively inhabit small streams spanning fewer than 100 square kilometers in Alabama, USA. One of these species, the Birmingham Darter *Etheostoma birminghamense*, was described in April 2025 from Valley Creek and its associated tributaries, which extend into the urban core of Birmingham, AL, and its suburbs. At least one population of *E. birminghamense* is feared extirpated, highlighting the imperilment of this microendemic species. Here, we report the results of recent collections that extend the range extension of *E. birminghamense* into Little Blue Creek, Nabors Branch, Halls Creek, and localities in the mainstem of Valley Creek. As previously hypothesized, occurrences of *E. birminghamense* are associated with exhumed Cambrian-Ordovician- and Mississippian-age carbonate units, which in Little Blue Creek appear only as pockets of exposed bedrock at the base of the channel and on the banks. Our observations demonstrate that *E. birminghamense* is distributed throughout the majority of the Valley Creek drainage, highlighting the need for rapid assessment of its conservation status.

## Introduction

Darters (*Percidae*: *Etheostomatinae*) are one of the most species-rich lineages of freshwater vertebrates inhabiting the southeastern North American freshwater biodiversity hotspot (Near et al. 2011; Page 1983). The diversity of this clade, which is endemic to North America, is asymmetrically distributed across this hotspot. The springs and creeks around the city of Birmingham in the US state of Alabama harbor a number of endemic species with very small ranges. These include the Federally Listed Rush Darter *Etheostoma phytophilum* Bart and Taylor 1999, Watercress Darter *Etheostoma nuchale* Howell and Caldwell 1965, and Vermillion Darter *Etheostoma chermocki* Boschung, Mayden and Tomelleri 1992. The Vermillion Darter is one of six species that comprise the *Etheostoma chermocki* species complex (Boschung et al. 1992; Clabaugh et al. 1996; Mayden and Kuhajda 2025; Brownstein et al. 2025; Kim et al. 2023).

Four of the six species of the *Etheostoma chermocki* species complex were recently delimited and described on the basis of a combination of genomic data and meristic and coloration traits (Brownstein et al. 2025; Kim et al. 2023; Mayden and Kuhajda 2025). These species, which last shared common ancestry more than one million years ago, face threats from stream degradation due to urbanization along with the Rush and Watercress Darters (Brownstein et al. 2025; Khudamrongsawat and Kuhajda 2007; Khudamrongsawat et al. 2005; Duncan et al. 2010; 2016; Fluker et al. 2010; 2009; Howell et al. 2016). The small ranges of these microendemic darter species and their proximity to urban centers makes updated and detailed documentation of their distribution essential (Howell et al. 2016; Brownstein et al. 2025). In the case of the *Etheostoma chermocki* complex, the close association of these species with carbonate bedrocks, which crop out only at specific sections of streams they occupy (Brownstein et al. 2025; Kim et al. 2023), suggests that even small habitat changes driven by anthropogenic activity could extirpate these ancient lineages or drive species to extinction.

Here, we report range extensions for the Birmingham Darter *Etheostoma birminghamense* Brownstein, Kim, Wood, Alley, Stokes, and Near 2025, which is endemic to the Valley Creek system around Birmingham and the adjacent city of Bessemer, Alabama (Brownstein et al. 2025). The Birmingham Darter population from Fivemile Creek has not been observed since 2006, underscoring its imperilment (Brownstein et al. 2025). We document the occurrence of the Birmingham Darter in Little Blue Creek, Halls Creek, and Nabors Branch, which fall within the lower, middle, and upper portions of the upper Valley Creek system. Although carbonate bedrock is not discernable from geological maps of some of the collection sites (Little Blue Creek), we documented its presence as these small streams erode through the capping layer of siliciclastic rock.

## Results and Discussion

We report *Etheostoma birminghamense* from six new localities all in Jefferson Co., Alabama: Little Blue Creek at CDX Gas Road crossing approximately 1.75 km southeast of Johns Road (AL 36), Bessemer; Valley Creek at shoal approximately 200 m downstream of Johns Road (AL 36) crossing, Bessemer; Valley Creek at B.Y. Williams Sr. Drive, Birmingham; unnamed tributary to Halls Creek near the Watercress Darter National Wildlife Refuge (NWR), approximately 60 m downstream of the culvert at the intersection of South Division Street and Division Court, Bessemer; Nabors Branch near 24th St SW, Birmingham, and in Seven Springs run (Nabors Branch tributary) along Cleburne Avenue, Birmingham (Figure 1).

**Figure 1.**
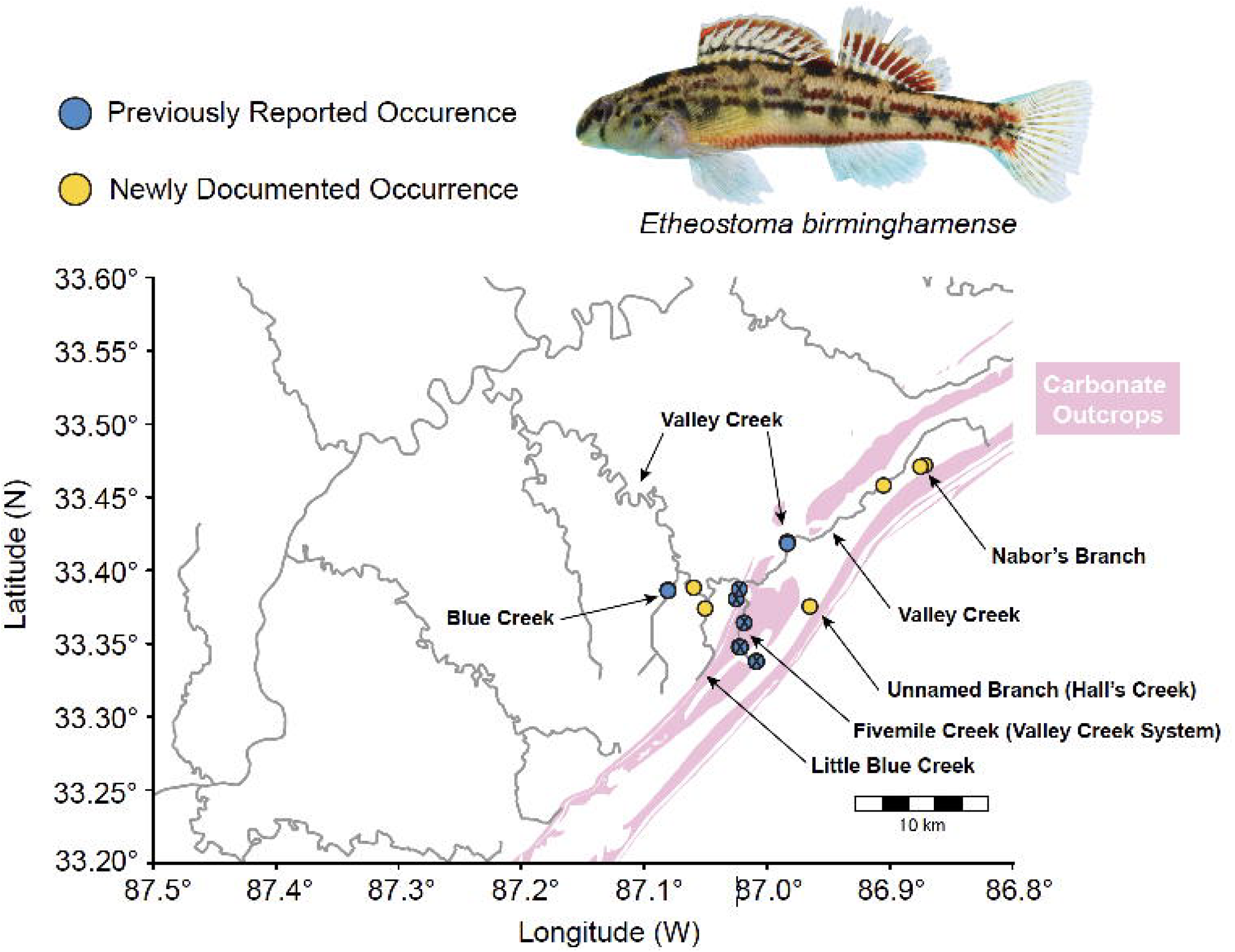
Locality map of Birmingham Darter *Etheostoma birminghamense*. Map shows rivers and streams in the Black Warrior River system and carbonate outcrops associated with them. Dots indicate localities from which *E. birminghamense* has been sampled. X marks on locality dots indicate extirpated populations. Photograph of Birmingham Darter by Julia E. Wood and Zachariah D. Alley, used with permission.

Biologists from Geological Survey of Alabama (including CBH, NDS) conducted a fish survey in an unnamed tributary to Halls Creek near the Watercress Darter NWR on December 4, 2024, during which three *Etheostoma birminghamense* [No voucher] were captured via seining approximately 60 m downstream of the culvert located at the intersection of South Division Street and Division Court. The three darters were originally identified as members of the *Etheostoma bellator* Suttkus and Bailey 1993, as *E. birminghamense* had not been formally described at the time. At the time of the survey, the current was slow, and the stream bed was primarily covered in coarse organic matter and detritus, with patches of gravel and exposed carbonate bedrock. Species that co-occur with *Etheostoma birminghamense* in the unnamed tributary to Halls Creek include Creek Chub *Semotilus atromaculatus*, Largescale Stoneroller *Campostoma oligolepis*, Blackspotted Topminnow *Fundulus olivaceus*, Western Mosquitofish *Gambusia affinis*, Bluegill *Lepomis macrochirus*, Gulf Longear Sunfish *Lepomis solis*, Alabama Bass *Micropterus henshalli*, Warrior Bass *Micropterus warriorensis*, Redspot Darter *Etheostoma artesiae*, Speckled Darter *Etheostoma stigmaeum*, and Watercress Darter *Etheostoma nuchale*.

Three of us (CDB, KA, TJN) collected three specimens [Yale Peabody Museum Ichthyology Collections (YPM ICH) XXXXX] of *Etheostoma birminghamense* from Little Blue Creek on October 11, 2025. We caught these specimens by seining along isolated patches of carbonate bedrock with moderate current on the stream margins. At this location, carbonate is barely exposed along the stream bed, which gravel and silt cover at its middle, while siliciclastic cap rock and soil emarginate its banks (Figure 2). Carbonate does not appear on geological maps of Cambrian-Ordovician and Mississippian outcrops of this stream (Figure 1) and is restricted to the margins of the stream bed.

**Figure 2.**
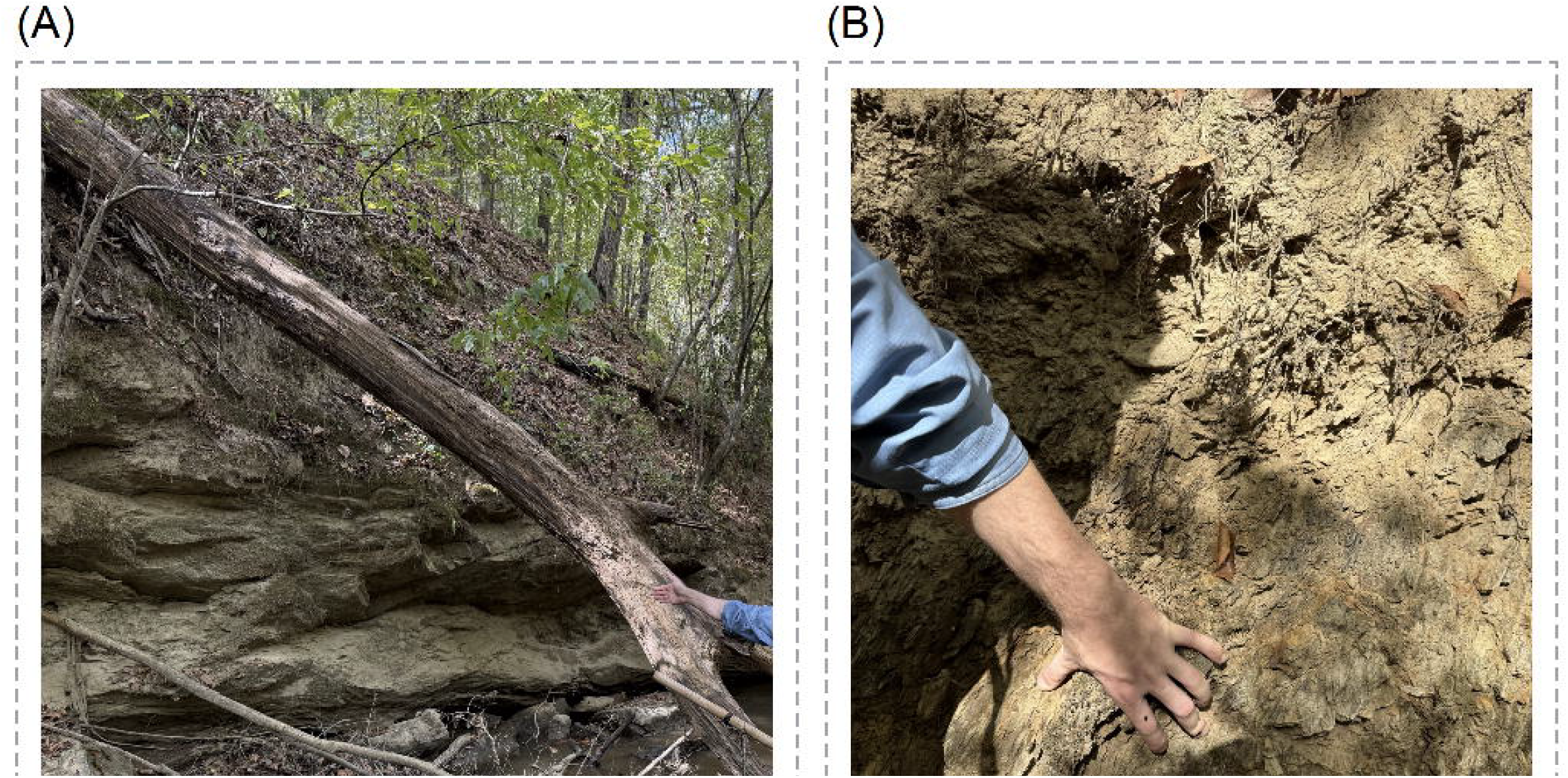
Examples of bedrock outcrops along Little Blue Creek. Photographs by KA, hand of CDB for scale. (A) Outcrops of overlying soil and siliciclastic rock on bank of Little Blue Creek, and (B) closeup of non-carbonate rock on side of stream.

Species that co-occur with *Etheostoma birminghamense* in Little Blue Creek include Blacktail Shiner *Cyprinella venusta*, Creek Chub *Semotilus atromaculatus, Campostoma oligolepis, Gambusia affinis, Lepomis macrochirus, Etheostoma artesiae*, and Blackbanded Darter *Percina nigrofasciata*. Upstream of this locality in Little Blue Creek off Mt. Forest Road, we collected numerous individuals of *E. artesiae*, as well as *Semotilus atromaculatus, Gambusia affinis*, and Gulf Longear Sunfish *Lepomis solis*, but did not observe the presence of *E. birminghamense*. Habitat at this upstream locality consisted of small, sporadically connected pools in the dry channel of Little Blue Creek.

In comparison, *Etheostoma birminghamense* is far more abundant in the main stem of Valley Creek approximately 200 m from Johns Road, Bessemer (18 individuals; YPM ICH XXXXX) and at B. Y. Williams Sr. Drive in Birmingham (64 individuals). At the Johns Road Valley Creek locality, *E. birminghamense* was abundant even though carbonate is not apparent on outcrop maps at this locality but was clearly present in the stream where it has been exhumed. We collected a small specimen of carbonate rock from this locality and is associated with the *E. birminghamense* specimen lot from this site in the YPM ichthyology collection. *Etheostoma birminghamense* appears especially abundant on carbonate that has been colonized by invertebrates and aquatic vegetation compared to uncolonized, slippery platforms. Here, co-occurring species include Alabama Hogsucker *Hypentelium etowanum* and *Percina nigrofasciata*.

On January 31, 2020, one of us (BRK) and colleagues collected over 40 *Etheostoma birminghamense* in Nabors Branch both above and below the mouth of Seven Springs run over gravel, cobble, and bedrock in slow to moderate flow. Other fishes present included *Campostoma oligolepis, Cyprinella venusta*, Silverstripe Shiner *Notropis stilbius, Semotilus atromaculatus, Hypentelium etowanum, Gambusia affinis, Lepomis solis, E. nuchale, E. stigmaeum*, and *Percina nigrofasciata*. This site was revisited on October 17, 2025 and 4 *E. birminghamense* were collected along with Banded Sculpin *Cottus carolinae*. Nabors Branch is the uppermost stream harboring *E. birminghamense* in the Valley Creek watershed and is a second stronghold for the species. Seven Springs is a small tributary to Nabors Branch, and on each of two dates (March 3, 2018 and October 17, 2025) a single *E. birminghamense* was collected over a silt and small gravel bottom with slow flow. Other fishes present included *Campostoma oligolepis*, Striped Shiner *Luxilus chrysocephalus, Semotilus atromaculatus, Hypentelium etowanum, Gambusia affinis*, and Green Sunfish *Lepomis cyanellus*.

On October 12, 2025, three of us (CDB, KA, TJN) captured only one individual of *Etheostoma birminghamense* (YPM ICH XXXXX) from the type locality of the species at Blue Creek, which was characterized by very low flow and moderate sedimentation. In Blue Creek, *E. birminghamense* co-occurs with *Hypentelium etowanum, Campostoma oligolepis*, Warrior Shiner *Lythrurus alegnotus*, Burrhead Shiner *Alburnops asperifrons, Notropis stilbius*, Speckled Darter *Etheostoma stigmaeum*, and *Percina nigrofasciata*.

The expansion of the known geographic distribution of the Birmingham Darter *Etheostoma birminghamense* reported in this manuscript suggests the species is widely distributed in the Valley Creek system in regions where there is sufficient water flow and outcropping carbonates. This supports the hypothesis that members of the *Etheostoma chermocki* species complex have speciated in a manner mediated by dispersal to areas in the upper Black Warrior River system with exposed carbonate bedrock (Kim et al. 2023). As previously noted, all members of this species complex face major risks to their survival from anthropogenic activity (Mayden and Kuhajda 2025; Brownstein et al. 2025). Specific threats to the Valley Creek watershed and *E. birminghamense* include an urban landscape (Brownstein et al. 2025; Mayden and Kuhajda 2025) with numerous impervious surfaces that result in high volumes of stormwater runoff and reduced recharge of groundwater, producing an unnatural hydrograph. This leads to streambed scouring after rain events, causing channelization and incision of stream beds and uprooting of aquatic vegetation, and exacerbated drought conditions in summer and fall further contributing to the loss of aquatic vegetation and resulting low ecological productivity. Springs within the Valley Creek watershed can mitigate low flow conditions but increased urban development and groundwater pumping put springs and their aquifers at risk (Duncan et al. 2010). The population of *E. birminghamense* in Fivemile Creek has not been observed for almost 20 years, despite a consistent documentation stretching from 1966 to 2006 and is feared extirpated (Brownstein et al. 2025) due to surface water disappearing into a sinkhole and pollution of the waterway from industrial discharge (“Black Warrior Riverkeeper - Press Releases” 2004).

We also highlight two specimen collections of single snubnose darters identified as *Etheostoma bellator* housed in the fish collection of the Auburn University Museum of Natural History fish collection (AUM). These specimens were collected in October 1971 from the Fivemile Creek system (Mayden and Kuhajda 2025), a direct tributary of Locust Fork—distinct from Fivemile Creek, which is a tributary of Valley Creek. During our October 2025 fieldwork, we attempted to collect snubnose darters from the same localities: AUM 9558 (Tarrant Spring Branch, Jefferson Co., Alabama 33.61333, -86.72639) and AUM 9624 (Fivemile Creek, Jefferson Co., Alabama 33.605, -86.73083). Despite extensive carbonate bedrock at the Fivemile Creek location, we failed to capture any snubnose darters. Other researchers have similarly failed to collect snubnose darters from Fivemile Creek over the last half century (W.M. Howell *pers. comm*. 2025), suggesting that this population, which may be distinct from all other species in the *Etheostoma chermocki* complex, is extinct. Adequate assessment of the status of Birmingham Darter in the Valley Creek system is essential for ensuring this brilliantly colored microendemic species does not suffer the same fate as this possibly distinct but undescribed species.

## Acknowledgements

We thank J.E. Wood, M.F. Stokes, B.P. Keck, D. Kim, and R.C. Harrington for discussions related to this manuscript. We thank Ms. Mary Rosenboom, Ms. Becky Morgan, Mr. David Havron, Mr. Marshall Killingsworth, and Lt. Col. (ret.) Ron Morgan for logistic support and warm hospitality. G. Watkins-Colwell provided collections support in the YPM Division of Vertebrate Zoology. We also thank the Freshwater Land Trust and U.S. Fish and Wildlife Service for logistical support.

## Permits

We confirm that we held all necessary permits for collections documented in this study, including a federal collections permit to BRK for sampling of Watercress Darter (TE22311A-4, TE22311A-5, ES22311A).

---

**Table 1.**
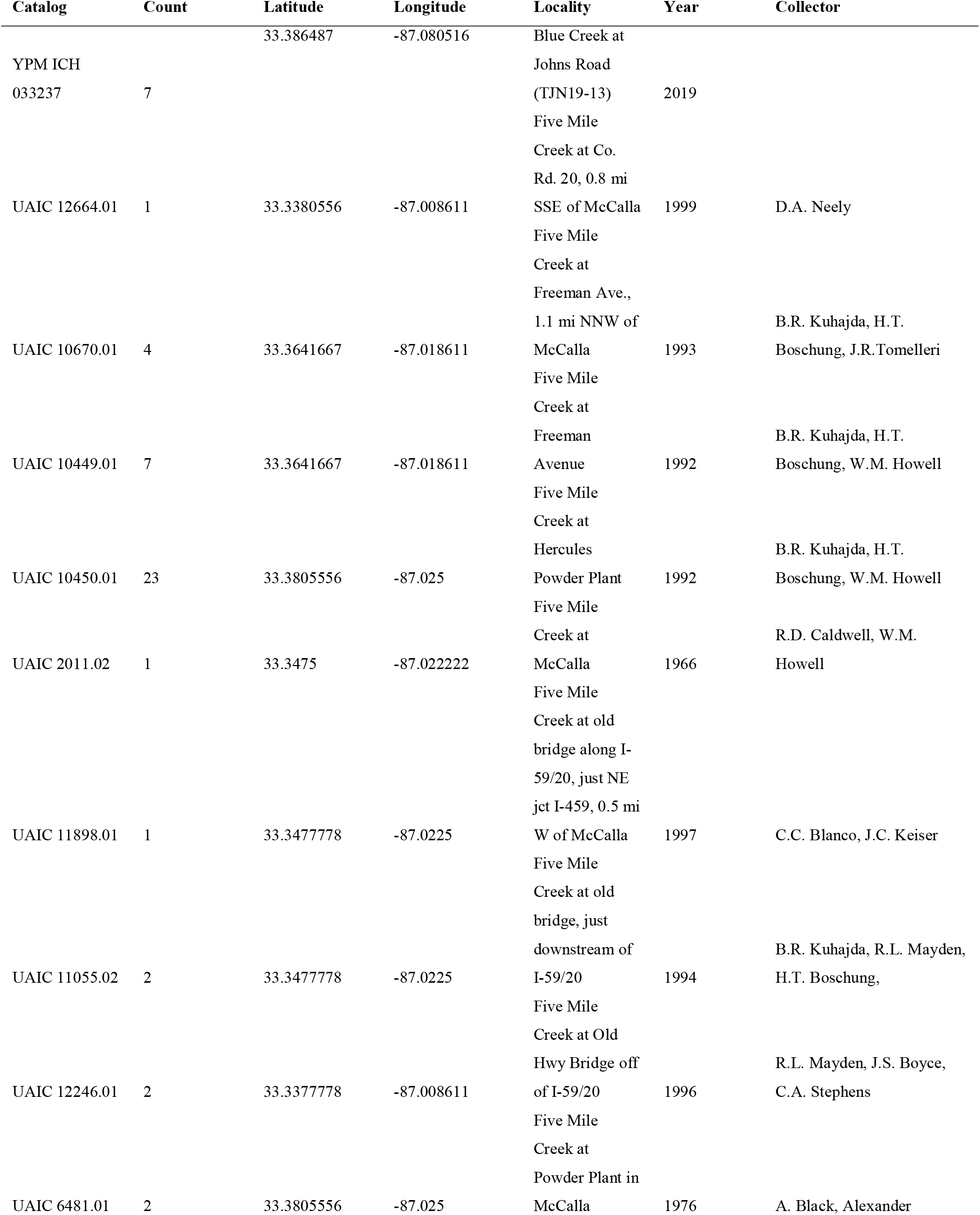
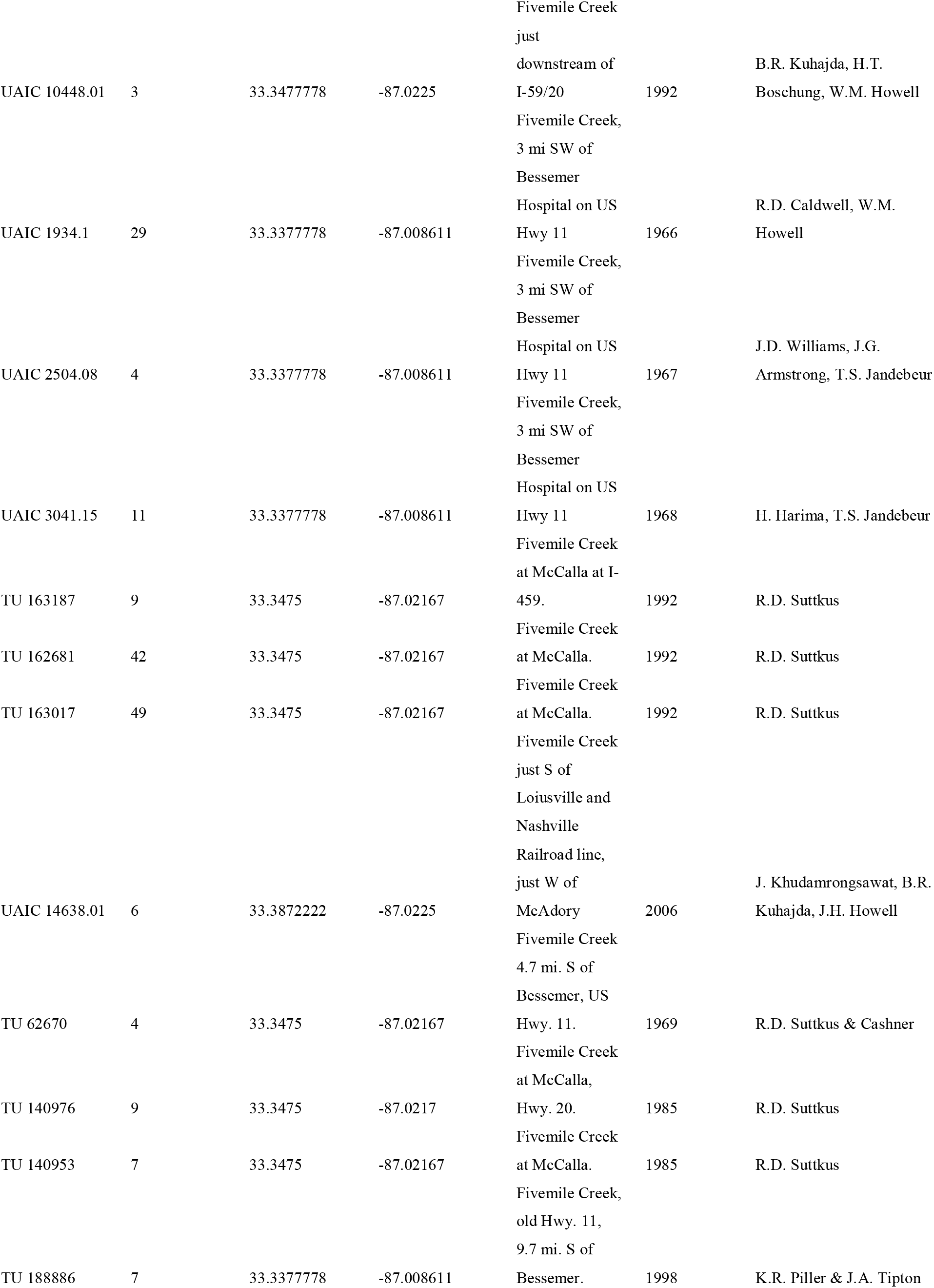
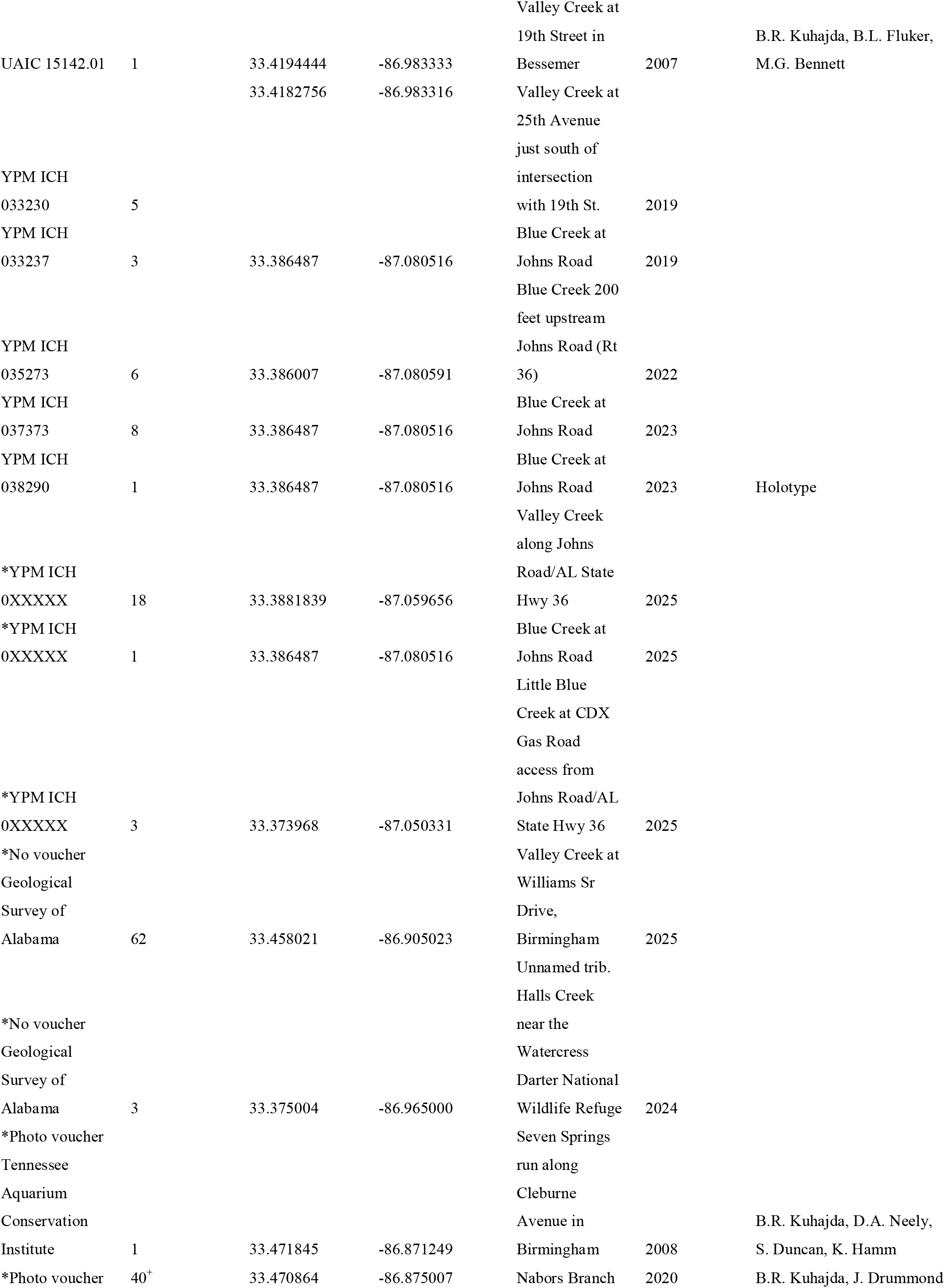
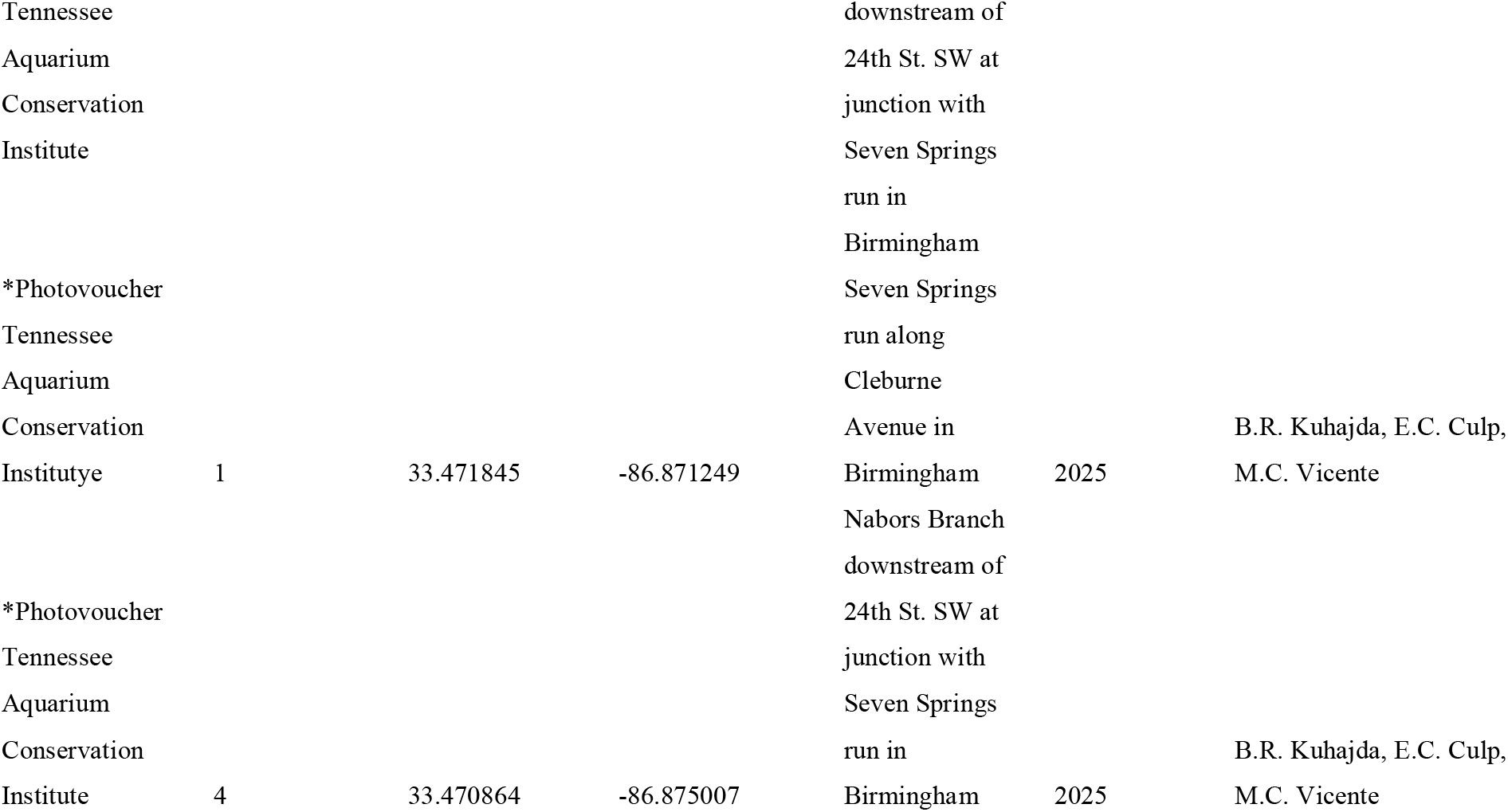
Full Set of Occurrences of Birmingham Darter (*Etheostoma birminghamense*) in Valley Creek watershed, Jefferson County, Alabama. New sites reported here denoted with asterisk.

